# Self-produced hydrogen sulfide improves ethanol fermentation by *Saccharomyces cerevisiae* and other yeast species

**DOI:** 10.1101/2022.08.09.503431

**Authors:** Emilio Espinoza-Simón, Paola Moreno-Alvarez, Elias Nieto-Zaragoza, Carolina Ricardez-García, Emmanuel Ríos-Castro, Salvador Uribe-Carvajal, Francisco Torres-Quiroz

## Abstract

Hydrogen sulfide (H_2_S) is a gas produced endogenously in organisms from the three domains of life. In mammals, it is involved in diverse physiological processes, including the regulation of blood pressure, and its effects on memory. In contrast, in unicellular organisms the physiological role of H_2_S has not been studied in detail. In yeast, for example, in the winemaking industry H_2_S is an undesirable byproduct because of its rotten egg smell; however, its biological relevance during fermentation is not well understood. The effect of H_2_S in cells is linked to a posttranslational modification in cysteine residues known as S-persulfidation. We evaluated S-persulfidation in the *Saccharomyces cerevisiae* proteome. We screened S-persulfidated proteins from cells growing in fermentable carbon sources and we identified several glycolytic enzymes as S-persulfidation targets. Pyruvate kinase, catalyzing the last irreversible step of glycolysis, increased its activity in the presence of a H_2_S donor. Yeast cells treated with H_2_S increased ethanol production; moreover, mutant cells that endogenously accumulated H_2_S produced more ethanol and ATP during the exponential growth phase. This mechanism of the regulation of the metabolism seems to be evolutionarily conserved in other yeast species, because H_2_S induces ethanol production in the pre-Whole Genome Duplication species *Kluyveromyces marxianus* and *Meyerozyma guilliermondii*. Our results suggest a new role of H_2_S in the regulation of the metabolism during fermentation.

## Introduction

Hydrogen sulfide (H_2_S) is a gasotransmitter produced endogenously in cells. It has been associated with diverse physiological processes such as vasodilation [1], pain [2] and longevity in animals [3], plant growth and development [4], bacterial antibiotic resistance [5], and as a byproduct of alcoholic fermentation in yeast [6]. Surprisingly, the biological function of H_2_S in yeast is not fully understood [7]; the majority of reports describe how it is produced or how to prevent its production during fermentation [8–10]. In yeast, H_2_S is involved in heavy metal detoxification [11], population synchrony [12], and chronological aging [3]; however the molecular mechanisms behind these phenomena have not been fully elucidated.

In yeast, the main metabolic pathway that produces H_2_S is the sulfate assimilation pathway [13], where inorganic sulfate is transformed to H_2_S and used in the synthesis of methionine and cysteine. This pathway is highly active in the exponential growth phase, as the principal H_2_S producer, sulfite reductase (encoded by *MET5* and *MET10)* [6], is highly active. The synthesis of H_2_S takes place in the first hours of fermentation and decreases at the final stages when cells reach the stationary phase [14]. The sulfur transferase Tum1p is another protein involved in H_2_S production during fermentation when high concentrations of cysteine are present in the media [15]. H_2_S is metabolized by Met17p a sulfhydrylase that catalyze the incorporation of sulfide for the biosynthesis of sulfur-containing amino acids [16].

The molecular effect of hydrogen sulfide depends on a posttranslational modification named S-persulfidation (originally termed sulfhydration) [17]. S-persulfidation involves addition of a thiol group to the cysteine residues (-S-SH) in proteins. This posttranslational modification has been associated with the activation and inhibition of protein activity [17,18].

In this work, for the first time, we evaluated the S-persulfidation of yeast proteins. We report that hydrogen sulfide is a regulator of glycolysis that increases ethanol production in *S. cerevisiae*. This was observed using an exogenous donor of hydrogen sulfide or mutant strains that accumulate or produce less H_2_S. This mechanism of regulation was conserved in pre-Whole Genome Duplication (WGD) species, such as the thermotolerant *Kluyveromyces marxianus* from the KLE clade and the oleaginous yeast *Meyerozyma guilliermondii* from the CUG-Ser1 clade. This work provides an insight into how H_2_S regulates glucose metabolism through an evolutionarily conserved mechanism, constituting an important role of H_2_S in fermentation.

## Materials and Methods

### Yeast strains, media, and growth conditions

*Saccharomyces cerevisiae* strains used in this study were S288C-derived laboratory strains BY4742 (*MATα his3Δ1 leu2Δ0 lys2Δ0 ura3Δ0*) referred as *wt*, BY4741 *(MATa his3Δ1 leu2Δ0 met17Δ0 ura3Δ0). Kluyveromyces marxianus* and *Meyerozyma guilliermondii* were isolated from mezcal producers in Michoacán, México [19]. Deletion strains derived from BY4742 were constructed by PCR-based gene replacement [20] using synthetic oligonucleotides and the kanMx and natMx disruption modules contained in plasmids pUG6 and pAG25. Gene deletions were confirmed by PCR using A and D oligos. Strains and oligonucleotides are listed in supplementary table 1. Strains were cultured at 30°C in liquid YPD medium (1% yeast extract, 2% dextrose, 2% peptone) or YPG medium (1% yeast extract, 2% galactose, 2% peptone) until reaching the exponential growth phase (optical density at 660nm [OD660] =0.5-0.6) and cells were collected then for protein extraction.

### Reagents

Sodium hydrosulfide (NaHS), Methyl methanethiosulfonate (MMTS), dithiothreitol (DTT), antibiotin antibody, neocuproine, deferoxamine and others chemicals were purchased from Sigma-Aldrich, St Louis MO, rabbit polyclonal anti-GAPDH (GTX100118, Genetex) were purchased from Genetex, Irvine, CA and N-(6-(biotinamido)hexyl)-3’-(2’-pyridyldithio)-propionamide (HPDP-biotin) (sc-207359), mouse monoclonal anti-enolase (sc-21738), goat polyclonal anti-CBS (sc-46830) and rabbit polyclonal anti-TIM (FL-249) were purchased from Santa Cruz Biotech, Dallas Tx.

### Modified biotin switch assay

The modified biotin switch assay was performed as described previously [17,21]. Briefly, after yeast cultures reached exponential phase, cells were collected, and intracellular proteins were extracted with chilled glass-beads in HEN buffer (250 mM HEPES-NaOH pH 7.7, 1 mM EDTA) supplemented with 1% triton X-100, 0.1 mM neocuproine, 0.1 mM deferoxamine and 1X protease cocktail inhibitor (Roche, Switzerland). Cell lysates were centrifuged at 16900 x *g* for 1 hr at 4°C, total extracts (1-2 mg) were blocked in HEN buffer with 2.5% SDS and 20 mM MMTS at 50°C for 20 min. The MMTS was removed by acetone precipitation and the protein pellet was resuspended in HEN buffer with 1% SDS. Protein labeling was performed with 0.8 mM HPDP-biotin for 3 h at room temperature in the dark. The biotinylated proteins were separated by SDS-polyacrylamide gel electrophoresis (PAGE) and subjected to immunoblot analysis.

### Purification of biotinylated proteins

After biotin switch assay, labeled extracts were subjected to streptavidin-based affinity precipitation with magnetic beads. Labeled extracts were incubated with 3X volumes of neutralization buffer (20 mM HEPES-NaOH pH 7.7, 100 mM NaCl, 1 mM EDTA, 0.5% triton) and 25 μl of streptavidin magnetic beads (Pierce) with agitation, overnight at 4°C. Magnetic beads were collected and washed with wash buffer as indicated by manufacturer’s instructions, biotinylated proteins were eluted with IP-MS elution buffer and analyzed using LS-MS or SDS-PAGE.

### Immunoblot analysis

Protein extracts were separated by SDS-PAGE and transferred to polyvinylidene difluoride (PVDF) membranes (Millipore-Merck, Germany). Membranes were blocked with 5% non-fat milk and incubated with a specific anti-biotin antibody overnight at 4°C. Proteins were detected with chemiluminescence using horser-adish peroxidase conjugated secondary antibodies (Jackson ImmunoResearch, West Grove, PA). Before to immunoblot, membranes were stained using Ponceau red (Millipore-Merck, Germany) as protein loading control.

### Quantification of intracellular ATP concentration

NaHS was added to cells cultures at specific timepoints, then cells were centrifuged, and intracellular ATP was measured using the ATP Bioluminescent Assay Kit HS II (Roche, Switzerland). Cell samples were prepared by diluting treated cells to a final concentration of 3.7×109 cells·mL^-1^ in 500 μl with a buffer containing 100 mM Tris-HCl pH 7.8 and 4 mM EDTA. After 2 min incubation, samples were immersed in boiling water for 2 minutes, and the resulting cell extracts were incubated for 5 minutes at 4°C, cell debris were removed by centrifugation at 16900 x *g* for 5 min and supernatants were used to measure the amount of intracellular ATP using an ATP calibration curve prepared each time, as indicated by the manufacturer. Bioluminescence was detected in a POLARstar Omega luminometer (BGM LABTECH, Offenburg, Germany). Three independent experiments with three replicas were performed, and values are represented as mean ± standard error.

### Detection of H_2_S production

H_2_S production by yeast strains colonies was detected through the generation of a visible black precipitate indicating that the hydrogen sulfide gas has reacted with lead nitrate [22]. Yeast strains were diluted, and cell density normalized to 3×10^7^ cells·mL^-1^. Cells were spotted in solid media (3.2% dextrose, 0.4% yeast extract, 0.24% peptone, 0.016% ammonium sulfate, 0.08% lead nitrate, 1.6% agar) and plates were kept at 30°C for 5-7 days. Also, H_2_S production was measured as reported previously [23] with some modifications. BY4742 *wt* strain were precultured at 30°C with constant shaking for 2 days in fresh YPD media. The assay was performed on a 96 well plate (COSTAR). Each well had 185 μL of YPD media, 5 μL of methylene blue (1 mg·mL^-1^) diluted in citrate buffer (100 mM, pH 4.5) and 10 μL of cells, for a final OD_600_ of 0.2. Growth was measured in an Infinite 200 (TECAN, Life Sciences) at 600 nm and 663 nm during 15 hours with intervals of 15 minutes between readings. During measures cells were incubated at 30°C with occasionally shaking. Three experimental replicates were made, with six different biological replicates in each experiment. Data for the hydrogen sulfide production were analyzed with the following formula:

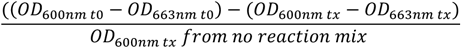

### Fermentation assays

BY4742 *wt* and derived mutants were precultured in liquid YPD medium for 24 h at 30°C, under agitation at 90 rpm in an Excella E24 incubator Shaker (New Brunswick Scientific, USA), then were inoculated in a 2L flask containing 500 mL of fresh YPD with an initial OD_600_=0.2 and incubated in the same conditions. When cells reached Do_660_=0.5, a pulse of NaHS was added. After 7h of NaHS addition, aliquots of 2 ml were obtained, DO_660_ was measured, and cells centrifuged at 16900 x *g* for 1 min. Supernatants were stored at −20°C for subsequent ethanol quantification. For mutant and *wt* strains, aliquots were taken every hour after cells were inoculated, supernatants were stored at −20°C. Ethanol production was evaluated through enzymatic assay coupled to NAD^+^ reduction. Briefly, supernatants were incubated in buffer (114 mM K_2_HPO_4_ pH 7.6), 1.8 mM NAD^+^ and 39μg·mL^-1^ alcohol dehydrogenase (ADH) for 30 min at 30°C with vigorous agitation [24]. Produced NADH was monitored by the increase in absorbance at 340 nm. The results are reported as mM ethanol per 1×10^7^ cells. Three independent experiments with three replicas were performed, and values are represented as mean ± standard error. *K. marxianus* fermentation assay was performed as in *S. cerevisiae* strains, when cells reached DO_660_=0.5, a pulse of NaHS 0.1mM was added. After 7h of NaHS addition, aliquots of 2 ml were obtained, and ethanol was qunatified. For *Meyerozyma guilliermondii* when cells reached DO_660_=0.5, a pulse of NaHS 0.1mM was added, 24h later another pulse of same concentration was added and 7h later ethanol was quantified.

### Enzymes activity assays

Glyceraldehyde 3 phosphate dehydrogenase (GAPDH) [17], pyruvate kinase (PK) [25] and alcohol dehydrogenase (ADH) [26] activities were measured by specific reaction assays and monitored spectrophotometrically at 340 nm, recording the rate of NAD to NADH reduction. Cells cultures were exposed to NaHS at different times, protein extracts were quantified, and 10 μg of protein were incubated in assay buffer as follows: for GAPDH (20 mM Tris-HCl pH 7.8, 100 mM NaCl, 0.1 mg·mL^-1^ Bovine serum albumin, 2 mM NAD^+^, 10 mM sodium pyrophosphate, 20 mM sodium arsenate, 500 mM DTT buffer, phosphate buffered saline (PBS) 1X, 27.3 mM glyceraldehyde 3-phosphate [G3P]). For PK (50 mM Imidazole·HCl, 120 mM KCl, 62 mM MgSO_4_ pH 7.6, 45 mM ADP, 6.6 mM NADH, 45 mM phosphoenolpyruvate [PEP], 1.3 KU·mL^-1^ lactate dehydrogenase). For ADH (114 mM K_2_HPO_4_ pH 7.6), 1.8 mM NAD+, and 16.4 mM ethanol). Three independent experiments with three replicas were performed, and values are represented as mean ± standard error.

### Oxygen consumption rate assay

BY4742 *wt* and derived mutant cells were precultured in liquid YPD medium for 48 hrs at 30°C, then were cultured in YPD medium with an initial DO_600_=0.2 under agitation in an Excella E24 incubator (New Brunswick Scientific, USA) for 7h at 30°C. Basal oxygen consumption was measured in resting cells with a Clark electrode (Oximeter model 782, Warner/Strathkelvin Instruments, North Lanarkshire, Scotland) in a water-jacketed chamber. Temperature was kept at 30 °C using a water bath (PolyScience 7 L, IL). Oxygen consumption reaction mixture was MES 10 mM pH 6 and 500 mg (wet weight) of cells were added at chamber [24]. To evaluate role of NaHS addition, when BY4742 cells reached OD_600_=0.5, a pulse of 0.1 mM NaHS was added. Seven hours later, basal oxygen consumption was measured as abovementioned. Results are reported as natgO/wet weight g/mn and values are represented as mean ± standard error. et weight) of cells were added at chamber.

### Sample preparation and LC-MALDI-MS/MS

Biotinylated proteins were digested with 250 ng of trypsin mass spectrometry grade (Sigma-Aldrich, St. Louis, MO) in 50 mM of ammonium bicarbonate (ABC). Resulting tryptic peptides were desalted using ZipTip C18 (Millipore) and concentrated to an approximated volume of 10 μL. Afterward, 9 μL were loaded into ChromXP Trap Column C18-CL precolumn (Eksigent, Redwood City CA); 350 μm X 0.5 mm, 120 A° pore size, 3 μm particle size and desalted with 0.1% trifluoroacetic acid (TFA) in H2O at a flow of 5 μL min^-1^ for 10 min. Then, peptides were loaded and separated on a 3C18-CL-120 column (Eksigent, Redwood City CA); 75 μm X 150 mm, 120 A° pore size, 3 μm particle size, in a HPLC Ekspert nanoLC 425 (Eksigent, Redwood City CA) using as a mobile phase A, 0.1% TFA in H2O and mobile phase B 0.1% TFA in acetonitrile (ACN) under the following lineal gradient: 0-3 min 10% B, 60 min 60% B, 61-64 min 90 % B, 65 to 90 min 10% B at a flow of 250 nL min^-1^. Eluted fractions were automatically mixed with a solution of 2 mg·mL^-1^ of alfa-cyano-4-hydroxycinnamic acid (CHCA) in 0.1% TFA and 50% ACN as a matrix, spotted in an Opti-TOF plate of 384 spots using a MALDI Ekspot (Eksigent, Redwood City CA) with a spotting velocity of 20 s per spot at a matrix flow of 1.6 μL min^-1^. The generated spots were analyzed by a MALDI-TOF/TOF 4800 Plus mass spectrometer (ABSciex, Framingham MA). Each MS Spectrum was acquired by an accumulation of 1000 shots in a mass range of 850-4000 Th with a laser intensity of 3800. The 100 more intense ions with a minimum signal-noise (S/N) of 20 were programmed to fragmenting. The MS/MS spectra were obtained by fragmentation of selected precursor ions using Collision-Induced Dissociation (CID) and acquired by 3000 shots with a laser intensity of 4300. Generated MS/MS spectrums were compared using Protein Pilot software v. 2.0.1 (ABSciex, Framingham MA) against *Saccharomyces cerevisiae*, strain ATCC 204508/S288c database (downloaded from Uniprot, 6049 protein sequences) using Paragon algorithm. Search parameters were: Not constant modifications in cysteines, trypsin as a cutter enzyme, all the biological modifications and amino acids substitution set by the algorithm (including carbamidomethylated cysteine as a variable modification); as well as phosphorylation emphasis and Gel-based ID as special factors. The detection threshold was considered in 1.3 to acquire 95% of confidence; additionally, the identified proteins showed a local FDR of 5% or less. Since a peptide derived from a given fragmentation spectra may be shared among redundant proteins during database search, it is necessary group all competing proteins and report only the protein with more spectrometric evidence; for this reason, identified proteins were grouped by ProGroup algorithm contained in the software Protein Pilot to minimize redundancy.

## Results

### S-Persulfidation of yeast proteins growing on a fermentable carbon source

Protein S-persulfidation was detected using the modified biotin switch method [17]. In order to validate the method in yeast, we performed the assay in either a poor producer (*met5*Δ*met10*Δ) or an accumulator (*met17*Δ) strain of H_2_S and compared them to the *wt* (BY4742) (Figure 1A). Cells were grown using glucose as the carbon source, and at the exponential phase, when H_2_S is produced [6], the protein was extracted. S-Persulfidated proteins were accumulated in *met17*Δ in comparison to the *met5*Δ*met10*Δ strain and wt as expected (Figure 1B).

**Figure 1.**
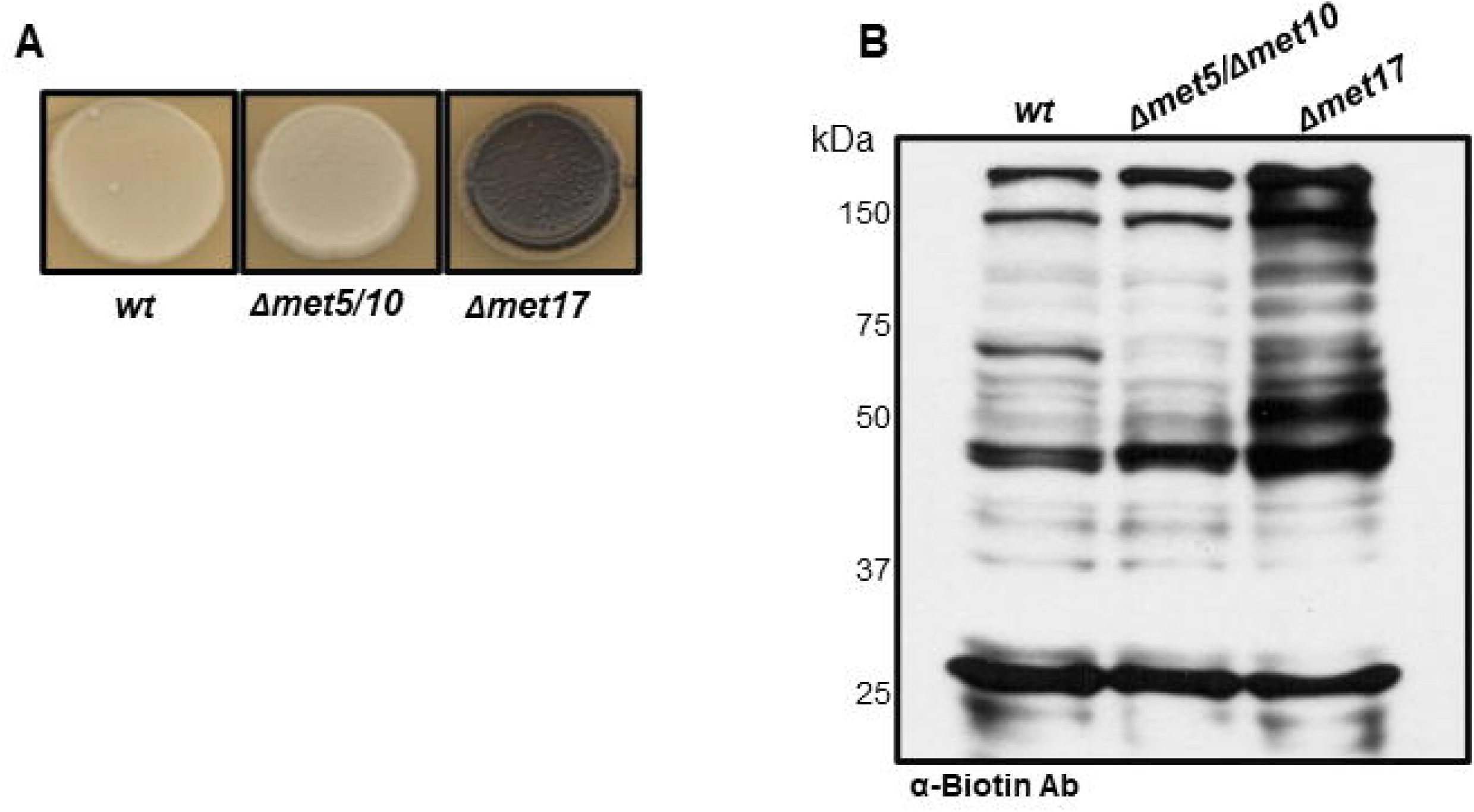
H_2_S productivity correlates with S-persulfidated proteins levels. (A) H_2_S productivity by *wt*, *met5*Δ*met10*Δ and *met17*Δ. Cells were incubated at 30°C on YPDL plates for 4 days. (B) S-persulfidated proteins in *wt*, *met5*Δ*met10*Δ and *met17*Δ strains. Whole cell extracts from exponential phase cultures were subjected to the modified biotin switch assay with antibody against biotin (α-Biotin Ab) to detect S-persulfidation.

In yeast, H_2_S is produced during fermentation, however, the S-persulfidation target proteins are not known. We used mass spectrometry to analyze the S-persulfidated proteins in cells growing at the exponential phase in two different fermentable carbon sources: glucose and galactose. Glucose is the preferred fermentable carbon source of yeast, while galactose needs to be isomerized to enter the glycolytic pathway. We found 42 S-persulfidated proteins; 21 were specific to glucose-grown cells, 4 were specific to galactose-grown cells, and 17 proteins were found in both conditions (Supplementary Table 2). Among the generally expressed 17 proteins, 15 were reported before as proteins with a redox-regulated cysteine [27], which is a feature of cysteines susceptible to posttrans-lational modifications [28]. Cytoplasmic translation (seven proteins) and glycolysis (seven proteins) where the most represented biological processes in the cells growing in either condition. Interestingly, pyruvate decarboxylase 1 (Pdc1), a key enzyme in alcoholic fermentation, and Adh1, the major enzyme responsible for ethanol synthesis, were also found, suggesting a possible role of S-persulfidation in fermentation. The identities of some glycolytic enzymes, glyceraldehyde 3-phosphate dehydrogenase (GAPDH), eno-lase, and triosephosphate isomerase (Tdh3, Eno2, and Tpi1, respectively) were confirmed using specific antibodies (Figure 2). We also tested cystathionine beta-synthase (Cys4), which was described before as a possible S-persulfidated protein [17, 29], although it must be considered that in our mass spectrometry analysis it did not pass the threshold (unused score of 1.04, coverage of 43.98%), we did find that Cys4 was a target of S-persulfidation.

**Figure 2.**
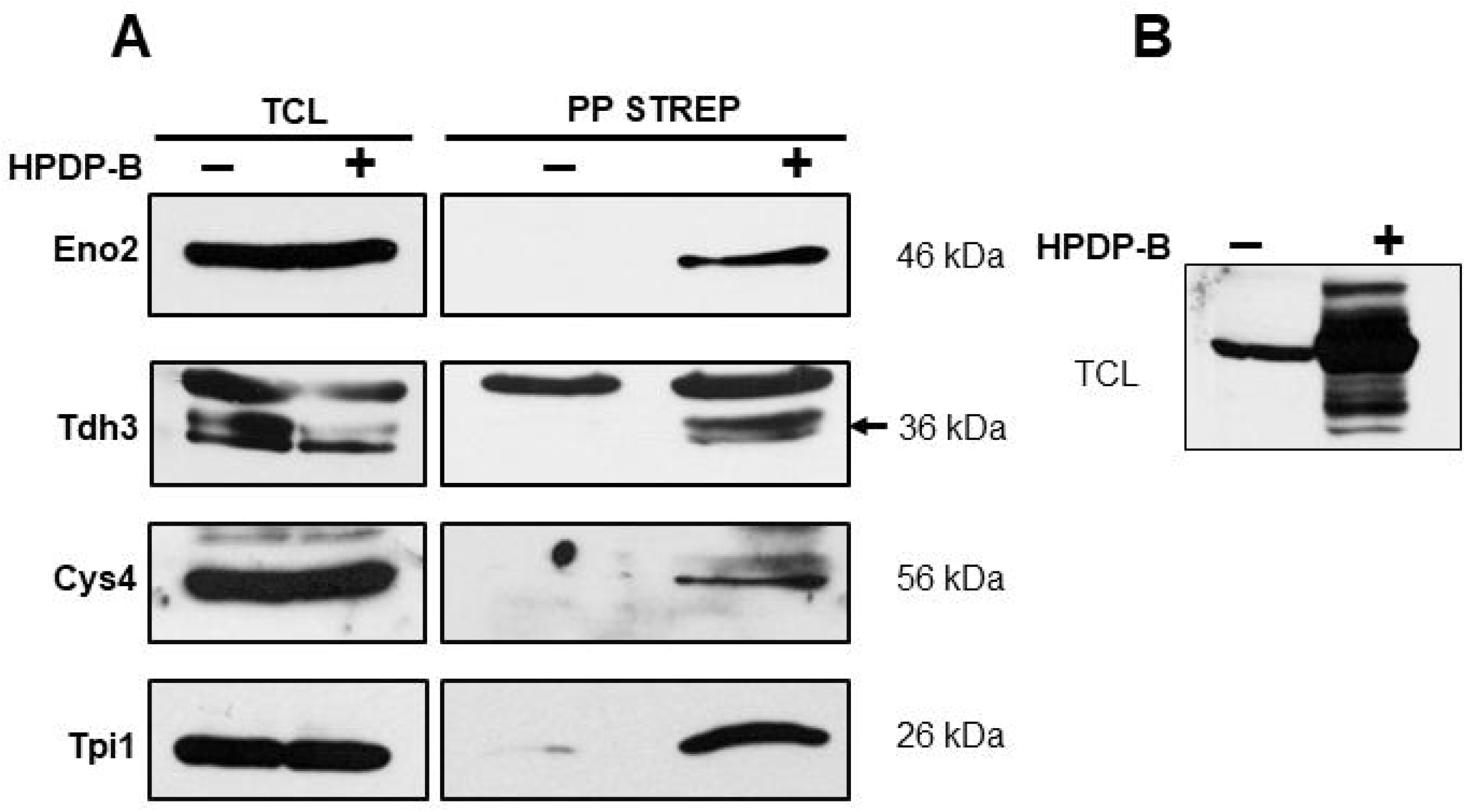
Confirmation of S-persulfidated proteins by streptavidin beads precipitation. A. Whole cell extracts from exponential phase cultures were subject to the modified biotin switch assay, precipitated with streptavidin beads and detected with antibodies specific to each protein; enolase (Eno2), glyceraldehyde 3 phosphate dehydrogenase (Tdh3), cystathionine beta synthase (Cys4) and triose phosphate isomerase (Tpi1). HPDP-B: (N-[6-(biotinamido)hexyl]-3’-(2’-pyridyldithio)propionamide). B. Whole cell extract from exponential phase cultures after modified biotin switch assay was used as input control. (-) line shows the proteins that reacts with anti-biotin antibody. (+) line shows biotinylated proteins after modified biotin switch assay. TCL: Total cell lysate, PP STREP: Streptavidin precipitation.

### H_2_S production during yeast growth

In order to evaluate H_2_S production during yeast growth, we determined H_2_S in the wt. The H_2_S reached a maximal amount during the log phase and dropped its production (Figure 3). Suggesting that the H_2_S concentration is not constant and drops when the culture stops growing.

**Figure 3.**
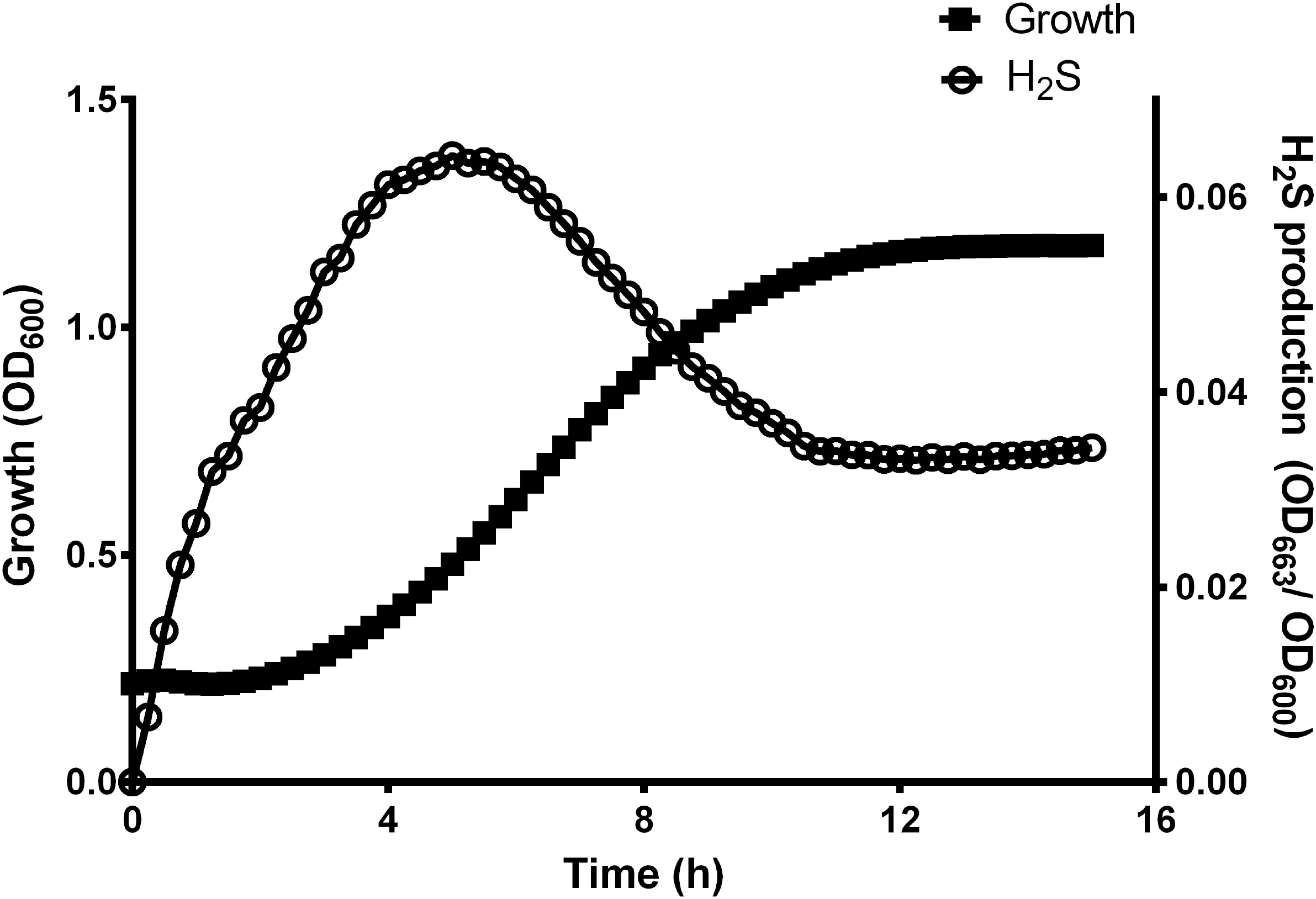
H_2_S production reach a maximal during log phase. Yeast cell were cultured in 96 wells microplate at 30°C and growth was measured at 600 nm. H_2_S was detected measuring methylene blue reduction at 663 nm.

### H_2_S increases glycolytic enzymes activities

Among the first effects of S-persulfidation described was the increase in GAPDH activity [17]. Considering that Tdh3 (GAPDH) was one of the glycolytic enzymes targeted for S-persulfidation, we decided to test the effect of NaHS (a donor of H_2_S) on GAPDH activity. Cells were stimulated with 0.1 mM or 0.25 mM NaHS for two and seven hours. Then, the protein was extracted, and the GAPDH activity was measured. We found that after two hours of NaHS stimulation, both at 0.1 mM and at 0.25 mM, increased GAPDH activity 1.4 times (Figure 4A), and the effect was lost at seven hours (Figure 4B). Another protein identified by mass spectrometry was pyruvate kinase (PK) that catalyzes the last irreversible step of glycolysis. We measured the activity of the pyruvate kinase at two hours of treatment with 0.1 mM or 0.25 mM NaHS finding that NaHS increased PK activity 2.39 times (Figure 4C, Supplementary Table 3), and lost its effect at seven hours (Figure 4D). Finally, we subjected alcohol dehydrogenase (ADH) to the same treatment, and we did not find any significant difference between the treated and untreated cells, i.e., at these concentrations NaHS did not affect ADH activity (Supplemental Figure 1).

**Figure 4.**
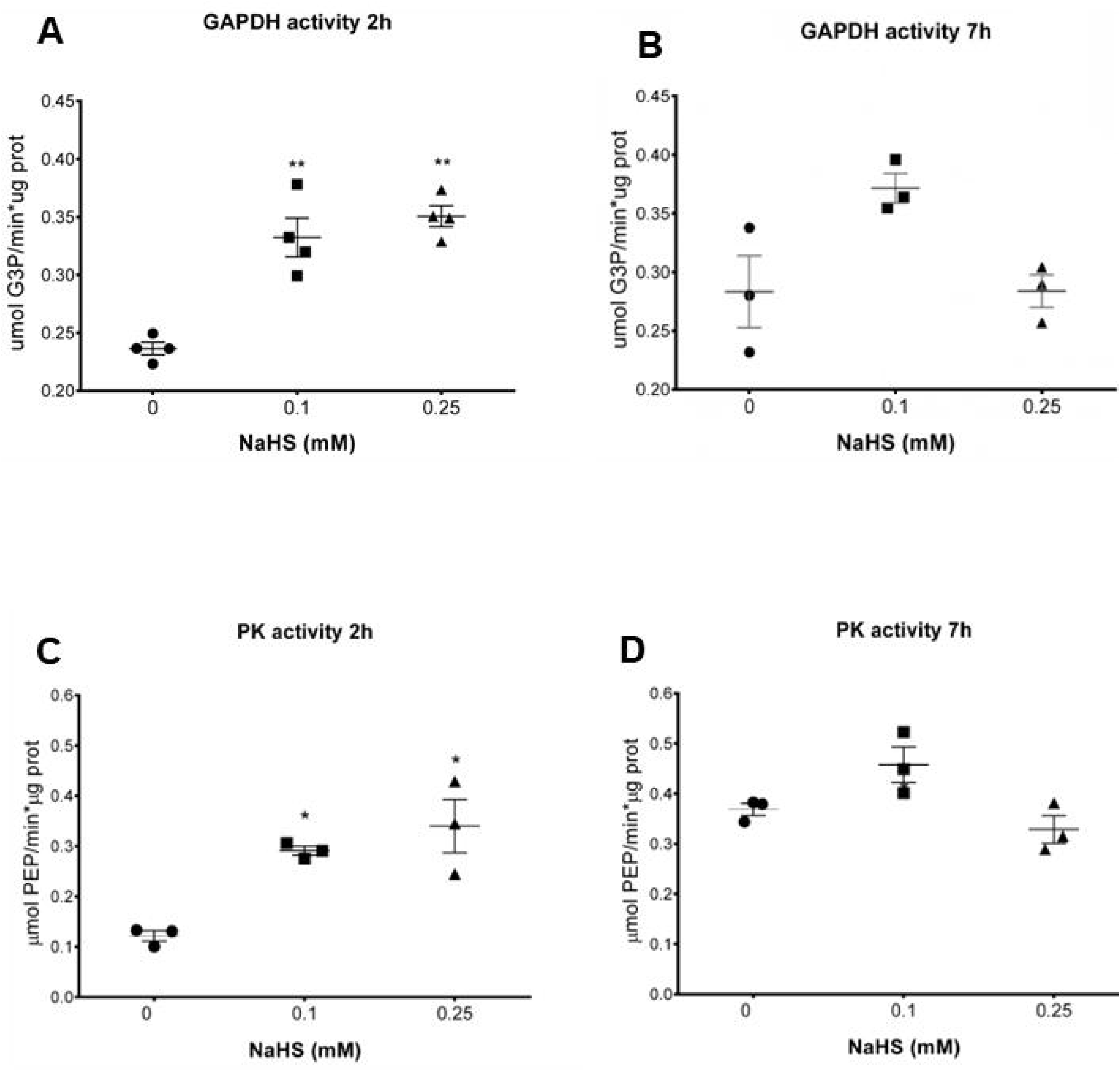
H2S increase the activity of GAPDH and Pyruvate Kinase two hours after stimu-lation. (A) Yeast cell cultures at exponential phase were treated with NaHS 0.1 and 0.25 mM. Two hours later whole cell extracts were used to measure GAPDH activity in vitro at 37°C. One-way ANOVA ** P<0.0001. (B) Yeast cell cultures at exponential phase were treated with NaHS 0.1 and 0.25 mM. Seven hours later whole cell extracts were used to measure GAPDH activity in vitro at 37°C. (C) Yeast cell cultures at exponential phase were treated with NaHS 0.1 and 0.25 mM. Two hours later whole cell extracts were used to measure Pyruvate Kinase activity in vitro at 37°C. (D) Yeast cell cultures at exponential phase were treated with NaHS 0.1 and 0.25 mM. Seven hours later whole cell extracts were used to measure Pyruvate Kinase activity in vitro at 37°C. One-way ANOVA * P<0.01. Closed circles, untreated cells; closed squares, NaHS 0.1 mM; closed triangles, NaHS 0.25.

### H_2_S stimulates fermentation

The glycolytic enzymes GAPDH and PK increased their activity in response to H_2_S. In addition, mass spectrometry data indicated that these and other enzymes from gly-colysis were S-persulfidated. Thus, we decided to test whether H_2_S influenced the syn-thesis of ethanol. Exponential-phase-grown cells were treated with 0.1 mM NaHS and after seven hours, the supernatant was collected. It was observed that the treated cells had increased ethanol production as compared to the control (Figure 5A).

**Figure 5.**
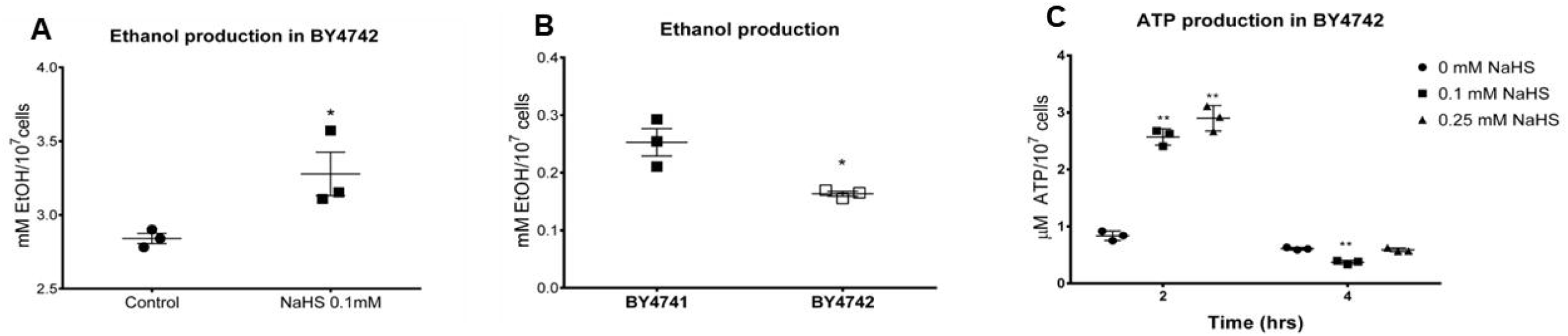
Exogenous and endogenous H_2_S on yeast cells induce ethanol production and ATP synthesis. (A) BY4742 yeast cell cultures were treated with NaHS 0.1 mM and seven hour later supernatants were collected. Ethanol production was measured in vitro at 37°C. Closed circles, untreated cells; closed squares, NaHS 0.1 mM. Unpaired t * P=0.04. (B) Yeast cell cultures of the strains BY4741 and BY4742 supernatants were collected at 24 h. Ethanol production was measured in vitro at 37°C. Closed squares, BY4741; open squares, BY4742. Unpaired t * P=0.02 (C) Yeast cell cultures at exponential phase were treated with NaHS 0.1 and 0.25 mM. Two and four hours later whole cell extracts were lysated and ATP was quantified. ATP production was measure in vitro at 37°C. Closed circles, untreated cells; closed squares, NaHS 0.1 mM; closed triangles, NaHS 0.25 mM. One-way ANOVA ** P<0.001.

In order to test the effect of endogenous H_2_S in fermentation, we decided to compare ethanol production in the two isogenic lab strains BY4741 and BY4742. The only difference between these two strains is that BY4741 has a deletion of *MET17* and strain BY4742 has a deletion of LYS2. As mentioned before, the *met17*Δ strain endogenously accumulates H_2_S, because *MET17* codifies for the enzyme using H_2_S and O-acetyl homoserine to synthetize homocysteine. An overnight preculture of each strain was diluted to an OD_600_=0.2, the supernatants were collected after 24 h, and the ethanol was quantified. The strain ac-cumulating H_2_S endogenously, BY4741 *met17*Δ, produced more ethanol than the strain BY4742 *MET17* (Figure 5B).

Ethanol is the main product of fermentation. Additionally, during glycolytic fermentation two ATP molecules are synthesized. Thus, we decided to quantify the ATP after treatment with NaHS. Exponential-phase cells were treated with the same quantities of NaHS used before, and the ATP was quantified two and four hours after treatment. After two hours, the treated cells produced more ATP than the untreated cells; this effect was lost four hours after treatment (Figure 5C). These results suggest that H_2_S stimulates glycolysis, and that fermentation is enhanced to produce both ATP and ethanol.

Based on these results we decided to compare whether the endogenous H_2_S had an influence on the onset of ethanol synthesis. We measured the ethanol production of the poor H_2_S producer strain *met5*Δ*met10*Δ, the H_2_S accumulator *met17*Δ strain, and the wt. The cells from 48 hours preculture were resuspended in fresh media and aliquots from the supernatant were collected every hour. The *met17*Δ strain initiated ethanol production at five hours, the wt strain initiated production at six hours, and the *met5*Δ*met10*Δ initiated production after seven hours (Figure 6A). This result showed that the cells with high endogenously accumulated H_2_S began ethanol production before the cells with a lower H_2_S concentration.

**Figure 6.**
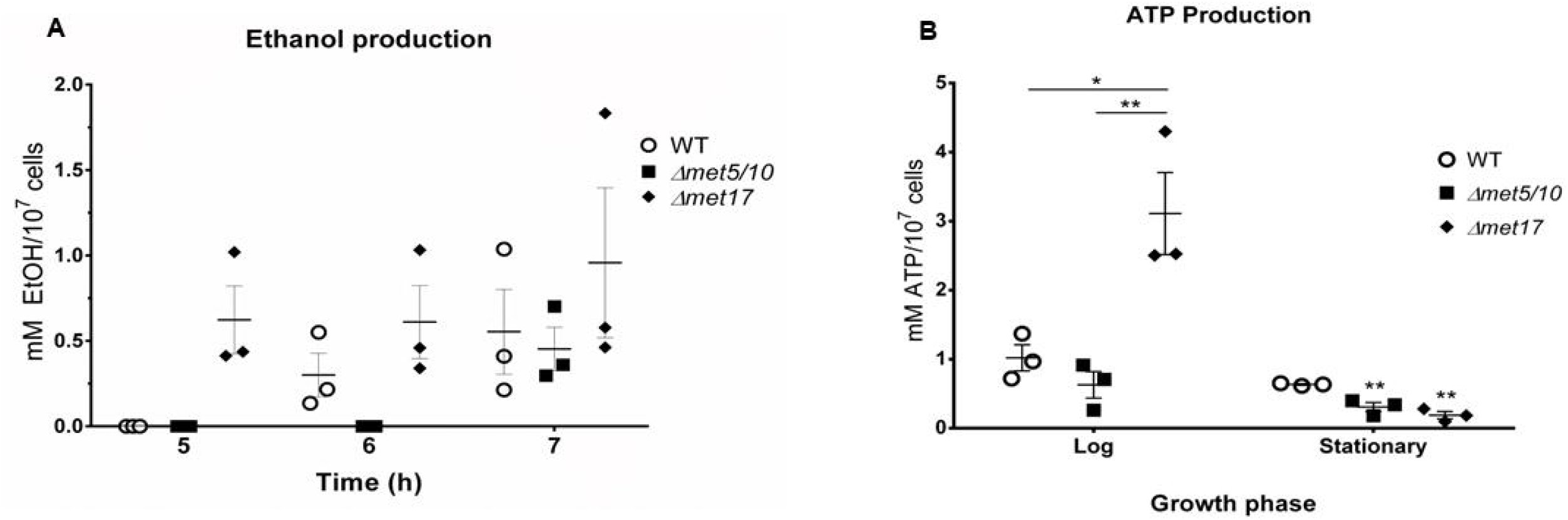
Yeast mutants that accumulate H_2_S synthesize ethanol faster and more ATP than lower endogenously accumulated H_2_S. (A) 48 hours precultures of yeast cell of the strains *wt*, *met5*Δ*met10*Δ and *met17*Δ were resuspended in fresh media and supernatants were collected every hour. Ethanol production was measured in vitro at 37°C. (B) Yeast cell cultures at exponential and stationary phase of the strains *wt*, *met5*Δ*met10*Δ and *met17*Δ were lysated and ATP was quantified. ATP production was measure in vitro at 37°C. One-way ANOVA * P<0.05, ** P<0.01. Open circles, *wt*; closed squares, *met5*Δ*met10*Δ; closed diamonds, *met17*Δ.

Finally, in each of these strains we quantified the ATP at the exponential or stationary phase. We found that the endogenously H_2_S accumulator strain produced the most ATP during the exponential phase, while there were no differences in the ATP concentration at the stationary phase between the wt and mutant strains (Figure 6B). These results support our proposal that H_2_S stimulates ethanol and ATP production.

### Endogenous H_2_S accumulation promotes basal oxygen consumption

ATP could be synthesized as product of the glycolysis and the oxidative phosphorylation. In order to elucidate if the ATP produced by H_2_S stimulation was from oxidative phosphorylation, we measured oxygen consumption from wt strain and mu-tants. A 48 hours preculture of each strain was diluted to an OD_600_= 0.2, and oxygen consumption was measured. Then, after seven hours (when cells were at exponential phase) oxygen was measured again (Figure 7). We found, in the diluted cells, that the *met5*Δ*met10*Δ, and the *met17*Δ consumed more oxygen than the *wt* strain. On the other hand, after seven hours of growing, we found that the *met17*Δ strain maintained the el-evated rate of oxygen consumption. This result suggest that endogenously accumulated H_2_S promotes oxygen consumption.

**Figure 7.**
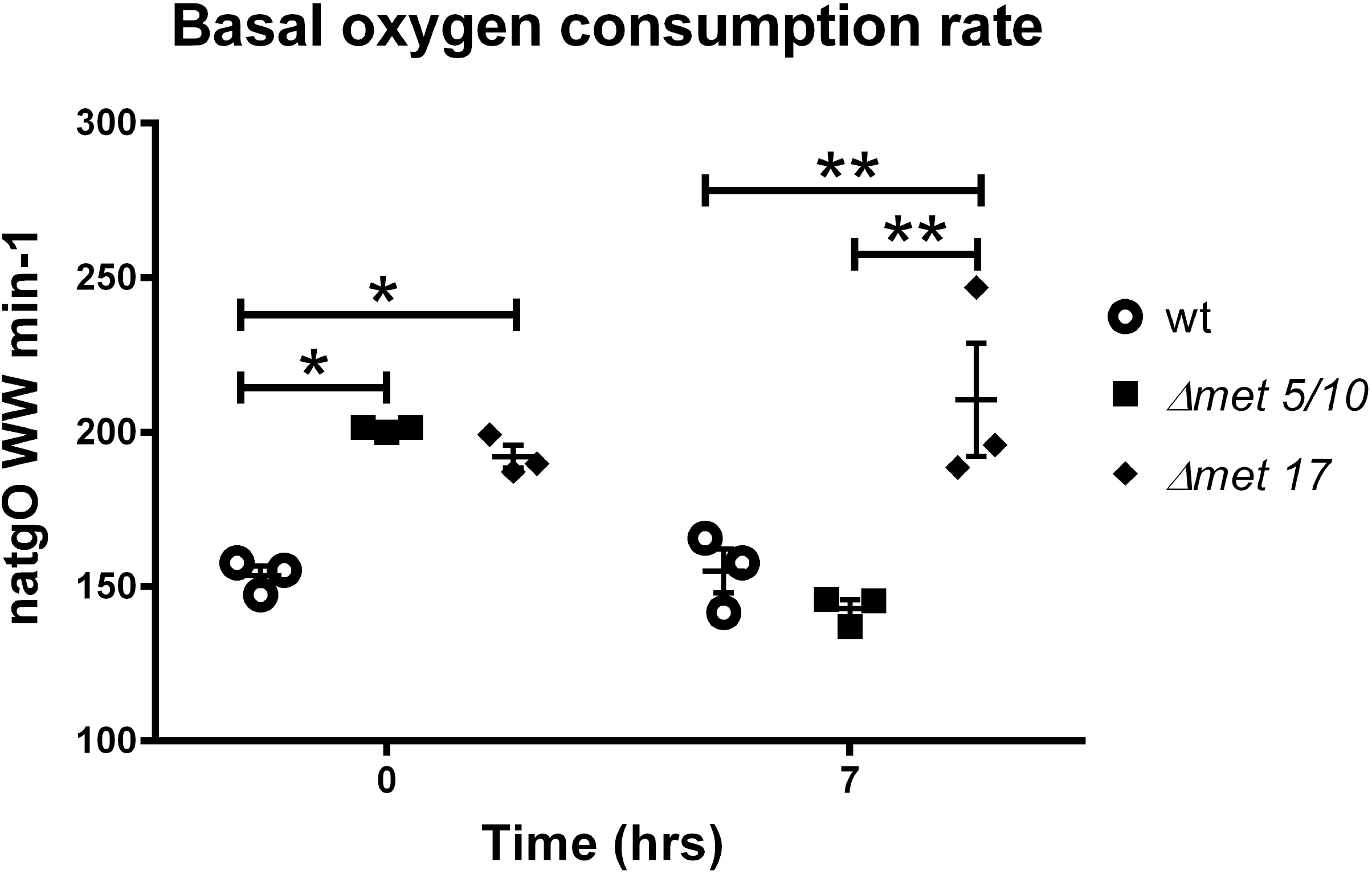
Endogenous H_2_S promotes basal oxygen consumption. 48 hours precultures of the *wt* and the mutants was diluted to an OD_600_= 0.2, and oxygen consumption was measured. The basal oxygen consumption was measured in resting cells in a Clark electrode at 30°C. Seven hours after dilution oxygen consumption was measured again. One-way ANOVA * P<0.05, ** P<0.01. Open circles, *wt*; closed squares, *met5*Δ*met10*Δ; closed diamonds, *met17*Δ.

### H_2_S stimulates ethanol production in Meyerozyma guilliermondii and Kluyveromyces marxianus

Ethanol synthesis is more robust in Crabtree positive yeast species; this phenomenon is associated with the WGD [30]. Evidence suggests that the WGD event arose from an interspecies hybridization between a strain from the KLE clade (genera Kluyveromyces, Lachancea and Eremothecium) and a strain from the ZT clade (Zygosaccharomyces and Torulaspora) [31]. The CUG-Ser1 clade first appeared approximately 117 million years before the WGD event; the CUG-Ser1 clade is characterized by a change in codon usage [32]. Considering this, we decided to test the effect of H_2_S during ethanol synthesis on the KLE clade strain, *K. marxianus* and in *M. guilliermondii* from the CUG-Ser1 clade. *K. marxianus* exponential-phase cells were stimulated with NaHS. Treatment with the H_2_S donor in *K. marxianus* increased ethanol synthesis as in *S. cerevisiae* (Figure 8a). In *M. guilliermondii*, it was noted that ethanol synthesis took longer than 24 h when glucose was the carbon source [33]; for this reason, the exponential-phase cells were treated with NaHS and treated again 24 h later. As observed in *K. marxianus*, ethanol synthesis increased after the H_2_S donor treatment on *M. guilliermondii* (Figure 8b) confirming that there is an effect of H_2_S on the fermentation activity of these two species of yeast.

**Figure 8.**
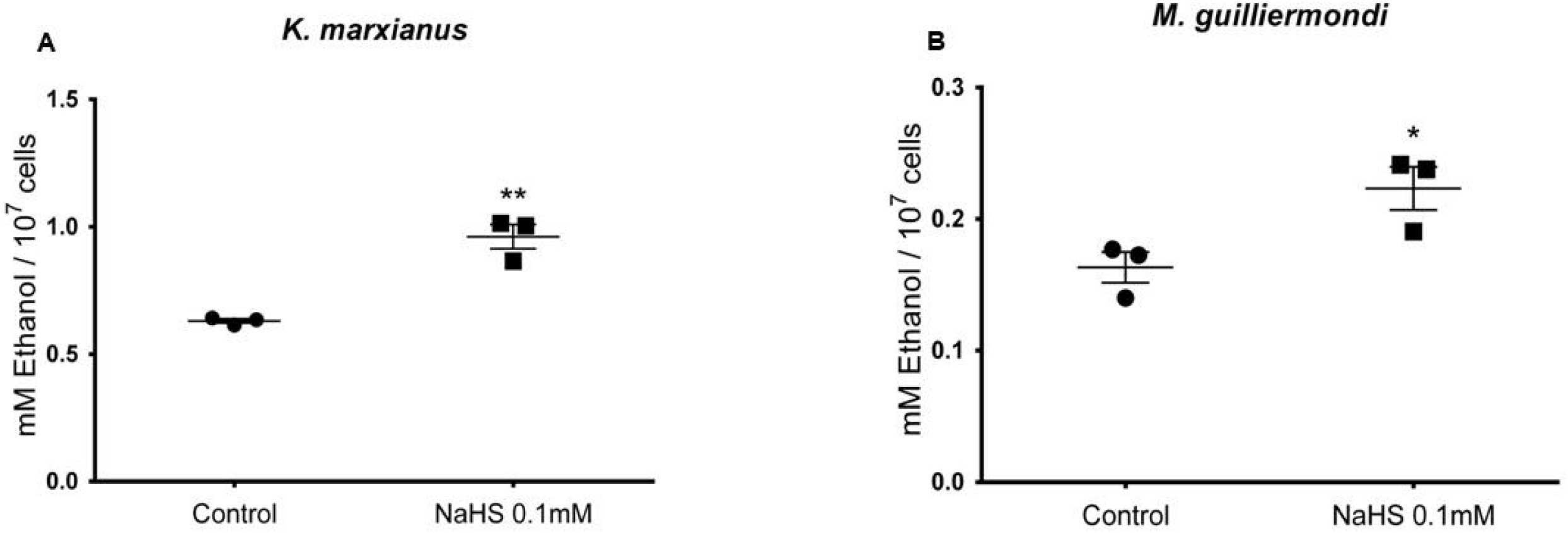
Exogenous H_2_S induce ethanol production in *K. marxianus* and *M. guilliermondi*. (A) *K. marxianus* yeast cell cultures were treated with NaHS 0.1 mM and seven hour later supernatants were collected. Ethanol production was measured in vitro at 37°C. Unpaired t ** P=0.002. (B) *M. guilliermondi* yeast cell cultures were treated with NaHS 0.1 mM and 24 h later cells were treated again with same concentration of NaHS. Seven hour later supernatants were collected. Ethanol production was measured in vitro at 37°C. Unpaired t * P=0.04. Closed circles, untreated cells; closed squares, NaHS 0.1 mM

## Discussion

Hydrogen sulfide is produced endogenously in yeast, and it is considered a fermentation byproduct; however its biological role is unknown. The biological effects of H_2_S are linked to a cysteine posttranslational modification termed S-persulfidation [17]. In this work, we analyzed S-persulfidated proteins on the yeast proteome. We identified several glycolytic enzymes as S-persulfidation targets, as reported previously in tissues such as the brain, heart and liver [29], in hepatocytes [17], a pancreatic beta cell line [34], HEK293 cells [29], plant [35,36] and bacteria [37]. Interestingly, Fu and collaborators reported six S-persulfidated glycolytic enzymes (ALDOA, GAPDH, PGK1, ENO1, PKM and LDHA) when they evaluated S-persulfidated proteins in cells overexpressing the H2S-producer enzyme cystathionine gamma-lyase (CSE) [29]. In a previous report, in pancreatic beta cells, metabolites were measured, and H_2_S was associated with an increased glycolytic metabolic flux of cells under chronic stress. All these reports agreed that several glycolytic enzymes were S-persulfidated, even when the activity of GAPDH was the only one measured [17]. Cysteine’s posttranslational modifications of glycolytic enzymes regulate the subcellular localization and oligomerization, which can impact its activity [38–40]. In a cell culture, H_2_S production is not constant, it starts to decline when the cells are at the middle of the logarithmic phase, suggesting that H_2_S synthesis could be regulated by the metabolic conditions. In *S. cerevisiae*, we found that GAPDH increased its activity with the H_2_S donor NaHS two hours after the treatment, and the effect was lost at seven hours. It is important to note that H_2_S is released from NaHS just a few seconds after the sodium salt is dissolved [41]; hence, the effect of NaHS two hours after treatment may be attributed to a chemical modification of the enzymes, such as S-persulfidation. The thioredoxin system eliminates this posttranslational modification [42,43], which is consistent with the idea that cellular mechanisms maintain protein S-persulfidation homeostasis [44]. Seven hours after stimulation, there was no effect on GAPDH activity probably due to the loss of S-persulfidation by the protein. Furthermore, the cells were no longer at an exponential phase seven hours after the OD_600_ reached 0.5, and the yeast metabolism changed to aerobic at the diauxic shift. The enzymes catalyzing the irreversible steps regulate the glycolytic pathway [45]; and we found that the enzyme involved in the last irreversible step, pyruvate kinase, increased its activity when stimulated with NaHS. The increase in pyruvate kinase activity may have at least two important consequences: feeding the Krebs cycle and/or stimulating the synthesis of ethanol. Considering that yeast synthe-tizes H_2_S during fermentation, we decided to test whether NaHS increased ethanol production. We found that cells treated with the H_2_S donor produced more ethanol. In order to confirm our observations, we measured the ethanol production in the isogenic strains BY4741 and BY4742. These strains have almost the same selection markers, and they differ only in one of them, BY4741 accumulates H_2_S because it is *met17*Δ; BY4742 is *lys2*Δ, and thus, it does not accumulate H_2_S. We observed that BY4741 produced more ethanol than BY4742, suggesting that endogenous H_2_S levels increase ethanol synthesis. Previously, it was reported that BY4742 fermenting activity was slower than in BY4741; it would be interesting to analyze the role of H_2_S in this system [46]. Considering these results, we proposed that if H_2_S stimulates fermentation, then mutants accumulating H_2_S would begin ethanol production before strains producing less H_2_S. We tested this hy-pothesis by comparing ethanol production between a lower producer of H_2_S, the strain *met5*Δ*met10*Δ, the accumulator strain *met17*Δ, and the *wt*. We found that the *met17*Δ strain started to produce ethanol before the wt strain; in turn, the *met5*Δ*met10*Δ strain production of ethanol was delayed even longer. The results confirmed that endogenous concentrations of H_2_S affects ethanol synthesis. Finally, we measured ATP production in all strains, and we found that at the logarithmic phase the *met17*Δ strain produced more ATP than the others. This result supports the idea that in addition to increasing an early synthesis of ethanol, H_2_S and also enhances ATP production at the exponential phase of growth.

The fermentation and the oxidative phosphorylation could yield ATP. We found that at exponential phase the *met17*Δ strain produced more ATP than the wt and *met5*Δ*met10*Δ strains. In order to test if the ATP was produced from the oxidative phosphorylation, we measured basal oxygen consumption on these strains. We found that the *met17*Δ strain has an elevated rate of oxygen consumption, and this is sustained when cells were at ex-ponential phase of growth; suggesting that endogenously accumulated H_2_S induces oxygen consumption. This would be contradictory to a report where described that ex-ogenous H_2_S inhibit respiration [47], however, at physiological concentrations H_2_S could induce the S-persulfidation of the ATP synthase from mammals and increases its activity [48]. The S-persulfidation takes place at cysteines 244 and 294 of human ATP synthase. The yeast ATP synthase (Atp1) conserved the cysteine 244 (Supplementary figure 2) in lineal sequence and has similar orientation on protein structure; therefore, the S-persulfidation could also be carried out in Atp1. Our results suggest that endogenous H_2_S has an effect on glycolysis and oxygen consumption. The effect of endogenous H_2_S on metabolism may explain the advantage of the *met17*Δ strain growing on fermentable carbon sources (glucose and galactose), over the *wt* and *met5*Δ*met10*Δ strains (Supple-mentary figure 3).

It is accepted that the origin of *S. cerevisiae* comes from a WGD event, probably by the interspecies hybridization between a strain from the KLE clade and a strain from the ZT clade [31]. WGD species have a more pronounced Crabtree effect than non-WGD species [30], so we decided to test whether H2S influenced yeast from the parental KLE clade that originated S. cerevisiae and a Crabtree-negative species from the CUG-Ser1 clade. The origin of this clade is estimated to be between 178 and 248 million years ago (mya), and this event occurred before WGD, estimated to be between 82 and 105 mya [32]. We found that both species increase ethanol production after the NaHS treatment suggesting that i) H2S is a positive regulator of fermentation and ii) this effect is evolutionarily conserved.

Overall, H2S is considered as a fermentation byproduct on yeast even when its biological effect is unknown. Here, we proposed a very different picture that will change our vision of how H2S regulates cell metabolism.

## 5. Conclusions

In conclusion, our data demonstrated that H_2_S is a regulator of energetic metabolism. These results fill a major gap in the understanding of H_2_S and its control of ethanol production, which is evolutionarily conserved among yeast species. Finally, our work provides the foundation for a mechanistic understanding of the effects of H_2_S.

## Supplementary Materials

Figure S1: Alcohol dehydrogenase activity; Figure S2: ATP synthase alignment; Figure S3: wt, met5Δmet10Δ, and met17Δ strains growth curves; Table S1: Strains and oligonucleotides primers; Table S2: Mass spectrometry results; Table S3: Protein activity data

## Acknowledgments

Authors acknowledge Dr. Antonio Peña and Dr. Norma Silvia Sanchez for the Kluyveromyces marxianus and Meyerozyma guilliermondii strains provided, Dr. Natalia Chi-quete-Félix, Dr. Gabriel del Rio and Dr. Teresa Lara for technical assistance.

## References

1. Yang, G.; Wu, L.; Jiang, B.; Yang, W.; Qi, J.; Cao, K.; Meng, Q.; Mustafa, A.K.; Mu, W.; Zhang, S.; et al. H2S as a Physiologic Vasorelaxant: Hypertension in Mice with Deletion of Cystathionine Gamma-Lyase. Science 2008, 322, 587–590, doi:10.1126/science.1162667.

2. Kawabata, A.; Ishiki, T.; Nagasawa, K.; Yoshida, S.; Maeda, Y.; Takahashi, T.; Sekiguchi, F.; Wada, T.; Ichida, S.; Nishikawa, H. Hydrogen Sulfide as a Novel Nociceptive Messenger. Pain 2007, 132, 74–81, doi:10.1016/j.pain.2007.01.026.

3. Hine, C.; Harputlugil, E.; Zhang, Y.; Ruckenstuhl, C.; Lee, B.C.; Brace, L.; Longchamp, A.; Treviño-Villarreal, J.H.; Mejia, P.; Ozaki, C.K.; et al. Endogenous Hydrogen Sulfide Production Is Essential for Dietary Restriction Benefits. Cell 2015, 160, 132–144, doi:10.1016/j.cell.2014.11.048.

4. Dooley, F.D.; Nair, S.P.; Ward, P.D. Increased Growth and Germination Success in Plants Following Hydrogen Sulfide Administration. PLoS One 2013, 8, e62048, doi:10.1371/journal.pone.0062048.

5. Shatalin, K.; Shatalina, E.; Mironov, A.; Nudler, E. H2S: A Universal Defense against Antibiotics in Bacteria. Science 2011, 334, 986–990, doi:10.1126/science.1209855.

6. Jiranek, V.; Langridge, P.; Henschke, P.A. Regulation of Hydrogen Sulfide Liberation in Wine-Producing Saccharomyces Cerevisiae Strains by Assimilable Nitrogen. Appl Environ Microbiol 1995, 61, 461–467, doi:10.1128/aem.61.2.461-467.1995.

7. Huang, C.-W.; Walker, M.E.; Fedrizzi, B.; Gardner, R.C.; Jiranek, V. Hydrogen Sulfide and Its Roles in Saccharomyces Cerevisiae in a Winemaking Context. FEMS Yeast Res 2017, 17, doi:10.1093/femsyr/fox058.

8. Huang, C.; Roncoroni, M.; Gardner, R.C. MET2 Affects Production of Hydrogen Sulfide during Wine Fermentation. Appl Microbiol Biotechnol 2014, 98, 7125–7135, doi:10.1007/s00253-014-5789-1.

9. Boudreau, T.F.; Peck, G.M.; O’Keefe, S.F.; Stewart, A.C. The Interactive Effect of Fungicide Residues and Yeast As-similable Nitrogen on Fermentation Kinetics and Hydrogen Sulfide Production during Cider Fermentation. J Sci Food Agric 2017, 97, 693–704, doi:10.1002/jsfa.8096.

10. Wang, C.; Liu, M.; Li, Y.; Zhang, Y.; Yao, M.; Qin, Y.; Liu, Y. Hydrogen Sulfide Synthesis in Native Saccharomyces Cerevisiae Strains during Alcoholic Fermentations. Food Microbiol 2018, 70, 206–213, doi:10.1016/j.fm.2017.10.006.

11. Sun, G.L.; Reynolds, Erin.E.; Belcher, A.M. Using Yeast to Sustainably Remediate and Extract Heavy Metals from Waste Waters. Nat Sustain 2020, 3, 303–311, doi:10.1038/s41893-020-0478-9.

12. Sohn, H.Y.; Murray, D.B.; Kuriyama, H. Ultradian Oscillation of Saccharomyces Cerevisiae during Aerobic Continuous Culture: Hydrogen Sulphide Mediates Population Synchrony. Yeast 2000, 16, 1185–1190, doi:10.1002/1097-0061(20000930)16:13<1185::AID-YEA619>3.0.CO;2-W.

13. Mendoza-Cózatl, D.; Loza-Tavera, H.; Hernández-Navarro, A.; Moreno-Sánchez, R. Sulfur Assimilation and Glutathione Metabolism under Cadmium Stress in Yeast, Protists and Plants. FEMS Microbiol Rev 2005, 29, 653–671, doi:10.1016/j.femsre.2004.09.004.

14. Oka, K.; Hayashi, T.; Matsumoto, N.; Yanase, H. Decrease in Hydrogen Sulfide Content during the Final Stage of Beer Fermentation Due to Involvement of Yeast and Not Carbon Dioxide Gas Purging. J Biosci Bioeng 2008, 106, 253–257, doi:10.1263/jbb.106.253.

15. Huang, C.-W.; Walker, M.E.; Fedrizzi, B.; Roncoroni, M.; Gardner, R.C.; Jiranek, V. The Yeast TUM1 Affects Pro-duction of Hydrogen Sulfide from Cysteine Treatment during Fermentation. FEMS Yeast Res 2016, 16, fow100, doi:10.1093/femsyr/fow100.

16. D’Andrea, R.; Surdin-Kerjan, Y.; Pure, G.; Cherest, H. Molecular Genetics of Met 17 and Met 25 Mutants of Saccharomyces Cerevisiae: Intragenic Complementation between Mutations of a Single Structural Gene. Mol Gen Genet 1987, 207, 165–170, doi:10.1007/BF00331505.

17. Mustafa, A.K.; Gadalla, M.M.; Sen, N.; Kim, S.; Mu, W.; Gazi, S.K.; Barrow, R.K.; Yang, G.; Wang, R.; Snyder, S.H. H2S Signals through Protein S-Sulfhydration. Sci Signal 2009, 2, ra72, doi:10.1126/scisignal.2000464.

18. Krishnan, N.; Fu, C.; Pappin, D.J.; Tonks, N.K. H2S-Induced Sulfhydration of the Phosphatase PTP1B and Its Role in the Endoplasmic Reticulum Stress Response. Sci Signal 2011, 4, ra86, doi:10.1126/scisignal.2002329.

19. Estrada-Ávila, A.K.; González-Hernández, J.C.; Calahorra, M.; Sánchez, N.S.; Peña, A. Xylose and Yeasts: A Story beyond Xylitol Production. Biochim Biophys Acta Gen Subj 2022, 1866, 130154, doi:10.1016/j.bbagen.2022.130154.

20. Bergkessel, M.; Guthrie, C.; Abelson, J. Yeast-Gene Replacement Using PCR Products. Methods Enzymol 2013, 533, 43–55, doi:10.1016/B978-0-12-420067-8.00005-2.

21. Paul, B.D.; Snyder, S.H. Protein Sulfhydration. Methods Enzymol 2015, 555, 79–90, doi:10.1016/bs.mie.2014.11.021.

22. Cost, G.J.; Boeke, J.D. A Useful Colony Colour Phenotype Associated with the Yeast Selectable/Counter-Selectable Marker MET15. Yeast 1996, 12, 939–941, doi:10.1002/(SICI)1097-0061(199608)12:10%3C939::AID-YEA988%3E3.0.CO;2-L.

23. Choi, K.-M.; Kim, S.; Kim, S.; Lee, H.M.; Kaya, A.; Chun, B.-H.; Lee, Y.K.; Park, T.-S.; Lee, C.-K.; Eyun, S.-I.; et al. Sulfate Assimilation Regulates Hydrogen Sulfide Production Independent of Lifespan and Reactive Oxygen Species under Methionine Restriction Condition in Yeast. Aging (Albany NY) 2019, 11, 4254–4273, doi:10.18632/aging.102050.

24. Peña, A.; Sánchez, N.S.; González-López, O.; Calahorra, M. Mechanisms Involved in the Inhibition of Glycolysis by Cyanide and Antimycin A in Candida Albicans and Its Reversal by Hydrogen Peroxide. A Common Feature in Candida Species. FEMS Yeast Res 2015, 15, fov083, doi:10.1093/femsyr/fov083.

25. Sakai, H.; Suzuki, K.; Imahori, K. Purification and Properties of Pyruvate Kinase from Bacillus Stearothermophilus. J Biochem 1986, 99, 1157–1167, doi:10.1093/oxfordjournals.jbchem.a135579.

26. Bergmeyer, H.V.; Gawhn, K.; Grassel, M. Alcohol Dehydrogenase. In Methods of Enzymatic Analysis; Academic Press: New York, 1974; Vol. 1, pp. 428–429.

27. Marino, S.M.; Li, Y.; Fomenko, D.E.; Agisheva, N.; Cerny, R.L.; Gladyshev, V.N. Characterization of Surface-Exposed Reactive Cysteine Residues in Saccharomyces Cerevisiae. Biochemistry 2010, 49, 7709–7721, doi:10.1021/bi100677a.

28. Alcock, L.J.; Perkins, M.V.; Chalker, J.M. Chemical Methods for Mapping Cysteine Oxidation. Chem Soc Rev 2018, 47, 231–268, doi:10.1039/c7cs00607a.

29. Fu, L.; Liu, K.; He, J.; Tian, C.; Yu, X.; Yang, J. Direct Proteomic Mapping of Cysteine Persulfidation. Antioxid Redox Signal 2020, 33, 1061–1076, doi:10.1089/ars.2019.7777.

30. Dashko, S.; Zhou, N.; Compagno, C.; Piškur, J. Why, When, and How Did Yeast Evolve Alcoholic Fermentation? FEMS Yeast Res 2014, 14, 826–832, doi:10.1111/1567-1364.12161.

31. Marcet-Houben, M.; Gabaldón, T. Beyond the Whole-Genome Duplication: Phylogenetic Evidence for an Ancient In-terspecies Hybridization in the Baker’s Yeast Lineage. PLoS Biol 2015, 13, e1002220, doi:10.1371/journal.pbio.1002220.

32. Shen, X.-X.; Opulente, D.A.; Kominek, J.; Zhou, X.; Steenwyk, J.L.; Buh, K.V.; Haase, M.A.B.; Wisecaver, J.H.; Wang, M.; Doering, D.T.; et al. Tempo and Mode of Genome Evolution in the Budding Yeast Subphylum. Cell 2018, 175, 1533–1545.e20, doi:10.1016/j.cell.2018.10.023.

33. Fabricio, M.F.; Valente, P.; Záchia Ayub, M.A. Oleaginous Yeast Meyerozyma Guilliermondii Shows Fermentative Metabolism of Sugars in the Biosynthesis of Ethanol and Converts Raw Glycerol and Cheese Whey Permeate into Polyunsaturated Fatty Acids. Biotechnol Prog 2019, 35, e2895, doi:10.1002/btpr.2895.

34. Gao, X.-H.; Krokowski, D.; Guan, B.-J.; Bederman, I.; Majumder, M.; Parisien, M.; Diatchenko, L.; Kabil, O.; Willard, B.; Banerjee, R.; et al. Quantitative H2S-Mediated Protein Sulfhydration Reveals Metabolic Reprogramming during the Integrated Stress Response. Elife 2015, 4, e10067, doi:10.7554/eLife.10067.

35. Aroca, Á.; Serna, A.; Gotor, C.; Romero, L.C. S-Sulfhydration: A Cysteine Posttranslational Modification in Plant Systems. Plant Physiol 2015, 168, 334–342, doi:10.1104/pp.15.00009.

36. Aroca, A.; Benito, J.M.; Gotor, C.; Romero, L.C. Persulfidation Proteome Reveals the Regulation of Protein Function by Hydrogen Sulfide in Diverse Biological Processes in Arabidopsis. J Exp Bot 2017, 68, 4915–4927, doi:10.1093/jxb/erx294.

37. Peng, H.; Zhang, Y.; Palmer, L.D.; Kehl-Fie, T.E.; Skaar, E.P.; Trinidad, J.C.; Giedroc, D.P. Hydrogen Sulfide and Reactive Sulfur Species Impact Proteome S-Sulfhydration and Global Virulence Regulation in Staphylococcus Aureus. ACS Infect Dis 2017, 3, 744–755, doi:10.1021/acsinfecdis.7b00090.

38. Aroca, A.; Schneider, M.; Scheibe, R.; Gotor, C.; Romero, L.C. Hydrogen Sulfide Regulates the Cytosolic/Nuclear Partitioning of Glyceraldehyde-3-Phosphate Dehydrogenase by Enhancing Its Nuclear Localization. Plant Cell Physiol 2017, 58, 983–992, doi:10.1093/pcp/pcx056.

39. Mitchell, A.R.; Yuan, M.; Morgan, H.P.; McNae, I.W.; Blackburn, E.A.; Le Bihan, T.; Homem, R.A.; Yu, M.; Loake, G.J.; Michels, P.A.; et al. Redox Regulation of Pyruvate Kinase M2 by Cysteine Oxidation and S-Nitrosation. Biochem J 2018, 475, 3275–3291, doi:10.1042/BCJ20180556.

40. Ford, A.E.; Denicourt, C.; Morano, K.A. Thiol Stress-Dependent Aggregation of the Glycolytic Enzyme Triose Phosphate Isomerase in Yeast and Human Cells. Mol Biol Cell 2019, 30, 554–565, doi:10.1091/mbc.E18-10-0616.

41. Li, L.; Whiteman, M.; Guan, Y.Y.; Neo, K.L.; Cheng, Y.; Lee, S.W.; Zhao, Y.; Baskar, R.; Tan, C.-H.; Moore, P.K. Characterization of a Novel, Water-Soluble Hydrogen Sulfide-Releasing Molecule (GYY4137): New Insights into the Biology of Hydrogen Sulfide. Circulation 2008, 117, 2351–2360, doi:10.1161/CIRCULATIONAHA.107.753467.

42. Ju, Y.; Wu, L.; Yang, G. Thioredoxin 1 Regulation of Protein S-Desulfhydration. Biochem Biophys Rep 2016, 5, 27–34, doi:10.1016/j.bbrep.2015.11.012.

43. Wedmann, R.; Onderka, C.; Wei, S.; Szijártó, I.A.; Miljkovic, J.L.; Mitrovic, A.; Lange, M.; Savitsky, S.; Yadav, P.K.; Torregrossa, R.; et al. Improved Tag-Switch Method Reveals That Thioredoxin Acts as Depersulfidase and Controls the Intracellular Levels of Protein Persulfidation. Chem Sci 2016, 7, 3414–3426, doi:10.1039/c5sc04818d.

44. Paul, B.D.; Filipovic, M.R. Editorial: Molecular Mechanisms of Thiol-Based Redox Homeostasis and Signaling in the Brain. Front Aging Neurosci 2021, 13, 771877, doi:10.3389/fnagi.2021.771877.

45. Campbell-Burk, S.L.; Shulman, R.G. High-Resolution NMR Studies of Saccharomyces Cerevisiae. Annu Rev Microbiol 1987, 41, 595–616, doi:10.1146/annurev.mi.41.100187.003115.

46. Harsch, M.J.; Lee, S.A.; Goddard, M.R.; Gardner, R.C. Optimized Fermentation of Grape Juice by Laboratory Strains of Saccharomyces Cerevisiae. FEMS Yeast Res 2010, 10, 72–82, doi:10.1111/j.1567-1364.2009.00580.x.

47. Wang, T.; Yang, Y.; Liu, M.; Liu, H.; Liu, H.; Xia, Y.; Xun, L. Elemental Sulfur Inhibits Yeast Growth via Producing Toxic Sulfide and Causing Disulfide Stress. Antioxidants (Basel) 2022, 11, 576, doi:10.3390/antiox11030576.

48. Módis, K.; Ju, Y.; Ahmad, A.; Untereiner, A.A.; Altaany, Z.; Wu, L.; Szabo, C.; Wang, R. S-Sulfhydration of ATP Synthase by Hydrogen Sulfide Stimulates Mitochondrial Bioenergetics. Pharmacol Res 2016, 113, 116–124, doi:10.1016/j.phrs.2016.08.023.

