## Supplemental Material for "Self-produced hydrogen sulfide improves ethanol fermentation by *Saccharomyces cerevisiae* and other yeast species"

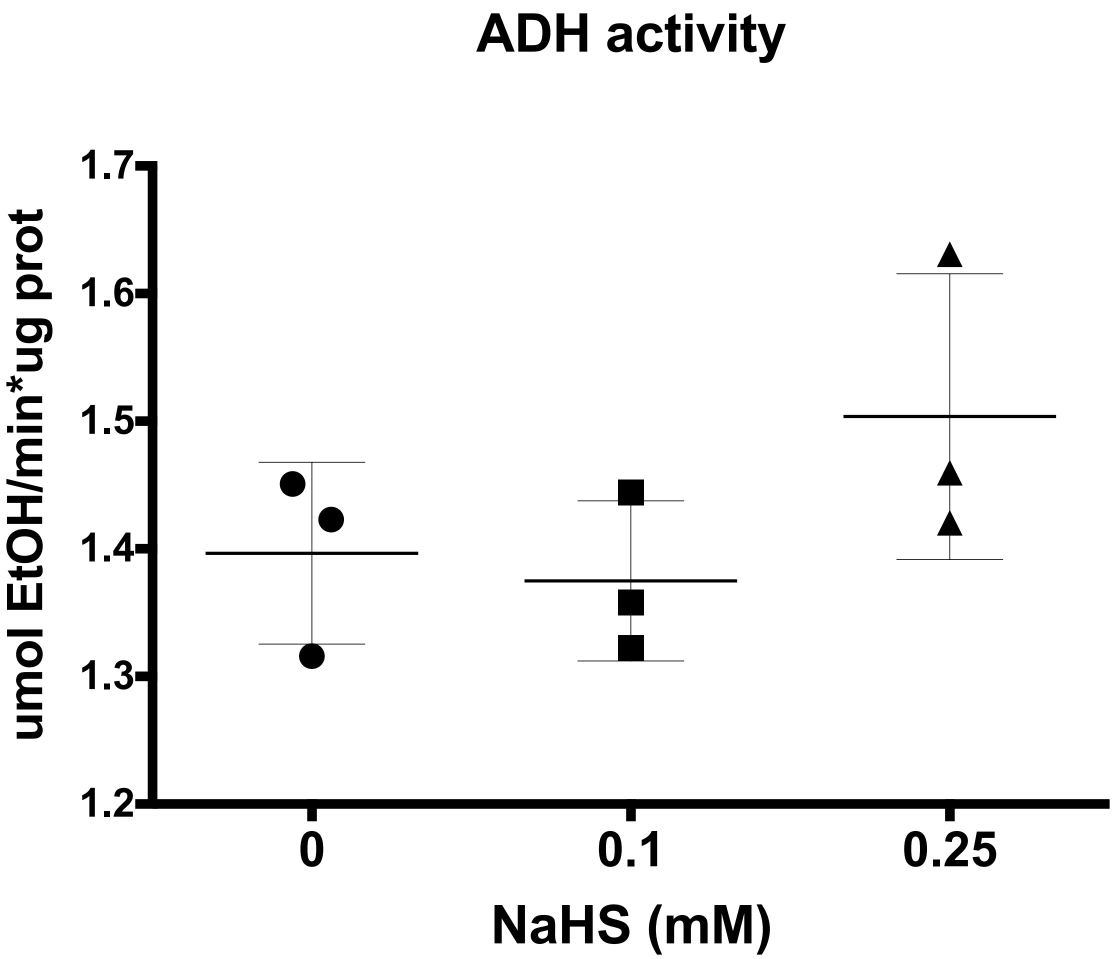


**Supplementary Figure 1. $H_2S$ does not increase the Alcohol dehydrogenase activity two hours after stimulation.** Yeast cell cultures at exponential phase were treated with NaHS 0.1 and 0.25 mM. Two hours later whole cell extracts were used to measure Alcohol dehydrogenase activity *in vitro* at 37°C.

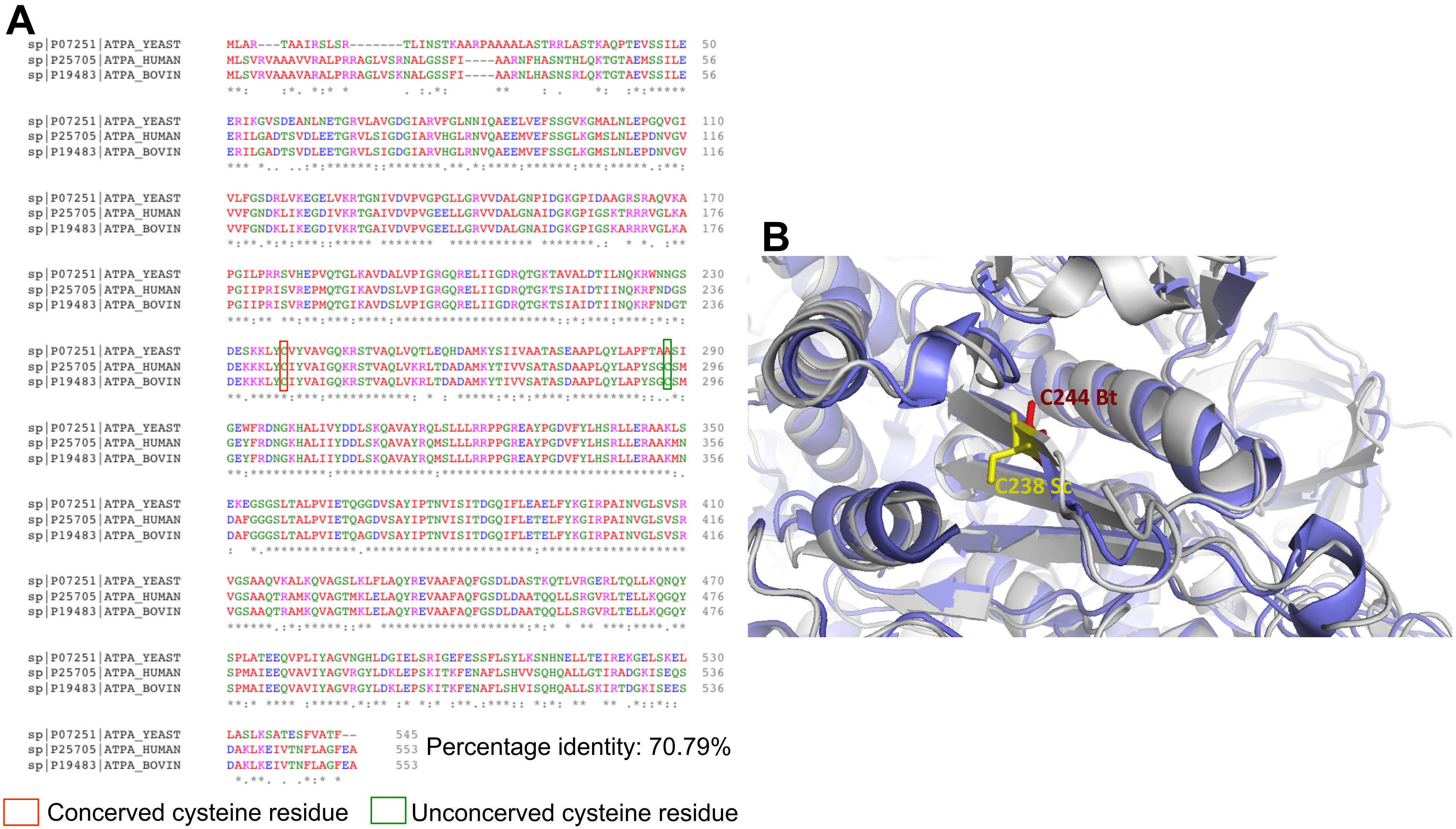


**Supplementary Figure S2.** ATP synthase cysteine residue conservation. (A) Sequence alignment of ATP synthases from *Saccharomyces cerevisiae, Homo sapiens* and *Bos taurus*. Mammal C244 and yeast C238 conserved residues are highlighted in red box. Mammal C294 residue is not conserved in yeast, also highlighted by green box. Lineal alignments were made using Clustal Omega. (B) Zoom from the structural superimposition of bovine ATP1A1 (blue, PDB code 5ARA) and yeast ATP5 (gray, PDB code 6CP6). Cysteine 238 from yeast (yellow, C238 Sc) and cysteine 244 from bovine (red, C244 Bt) are in the same sheet with similar orientation. Nearly all secondary structures from local vicinity are conserved between mammals and yeast. Structural superimposition were made using PyMOL.

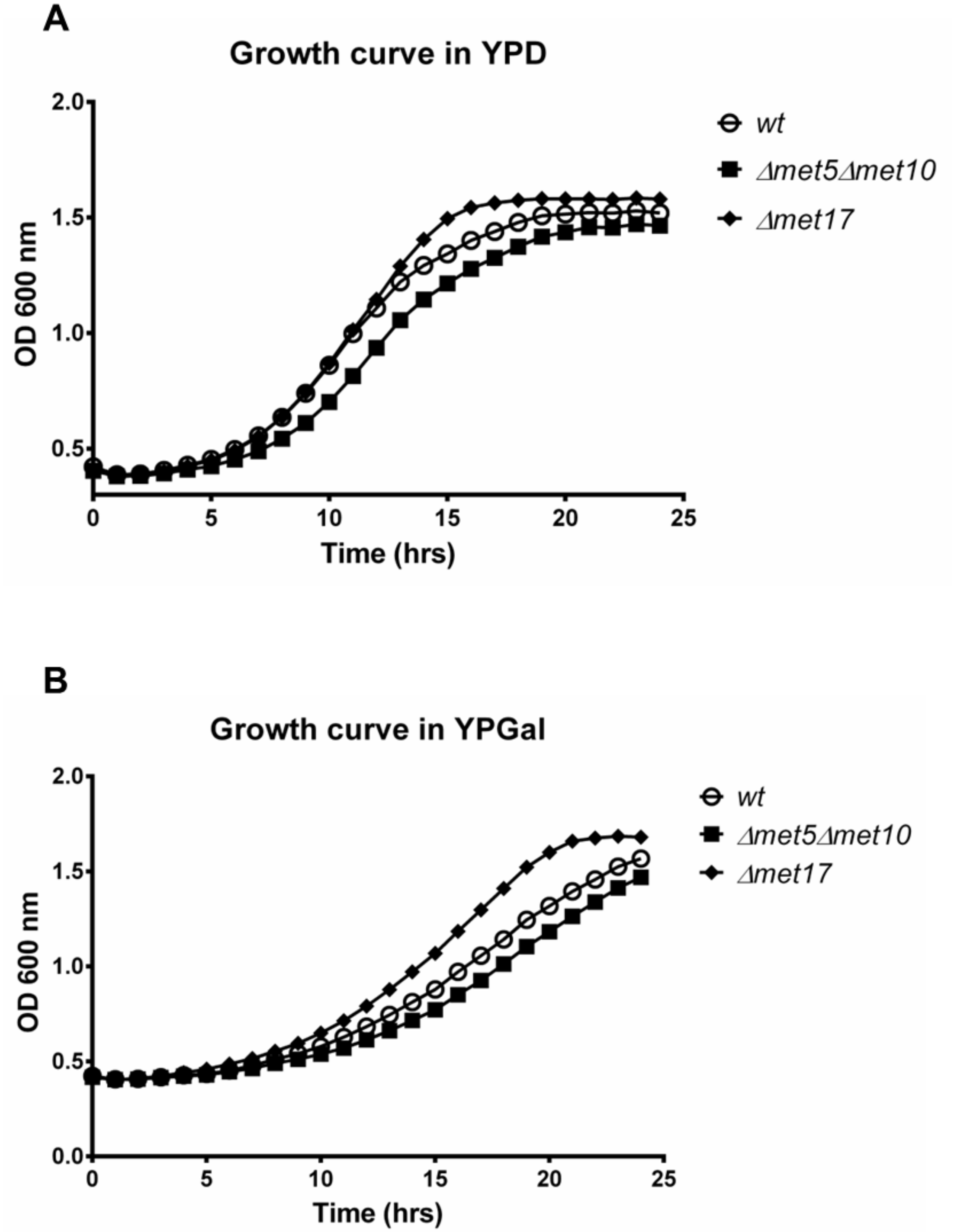


**Supplementary Figure 3. $H_2S$ overproducer strain grows faster on fermentable culture medium.** *S. cerevisiae* BY4742 (*wt*) and derived mutants were precultured in YPD medium for 24 h at 30°C under constant agitation. Next, cells were inoculated at $OD_{600}$=0.03 in A) YPD or B) YPGal for 24 hrs and $OD_{600}$ nm in a Bioscreen C microplate absorbance reader were measured.

### Supplementary Table 1

Yeast strains

| Name | Genotype | Source |
|---|---|---|
| BY4741 | *MAT**a**; his3Δ 1; leu2Δ 0; met17Δ 0; ura3Δ* | Euroscarf |
| BY4742 | *MATα; his3Δ 1; leu2Δ 0; lys2Δ 0; ura3Δ* | Euroscarf |
| GNO13 | *MATα; his3Δ 1; leu2Δ 0; lys2Δ 0; ura3Δ; met5Δ:KanMx; met10Δ:ClonNAT* | This study |
| PM01 | *MATα; his3Δ 1; leu2Δ 0; lys2Δ 0; ura3Δ; met17Δ:KanMx* | This study |
| *Kluyveromyces marxianus* | Isolated from mezcal production | Estrada-Ávila 2022 |
| *Meyerozyma guilliermondii* | Isolated from mezcal production | Estrada- Ávila 2022 |

Oligonucleotide primers

| Name | Secuence |
|---|---|
| met5F | ATAAAGTAACAGTAGGGAACGGGAACCAGAGAAAAACAAAAGATTGGCGAcagctgaagcttcgtacgc |
| met5R | AATGTGGGAAGAAAACCCAATAGTATGTCCTACTATGTCATATGCTATCAcataggccactagtggatc |
| met10F | ATAGTTTTTTTCAACTGCTTTTCCTCGAGGTCACCCAAATATACAACGAGcagctgaagcttcgtacgc |
| met10R | TTCAAGACAGGTTCAATAAATAGATATTTAGTTTTTATTACTATATTAATcataggccactagtggatc |
| met17F | TACAGGGTCGTCAGATACATAGATACAATTCTATTACCCCCATCCATACAcagctgaagcttcgtacgc |
| met17R | TTGTGAGAGAAAGTAGGTTTATACATAATTTTACAACTCATTACGCACACcataggccactagtggatc |
| met5A | TTCATCACGTGCGTATTATCTCTTA |
| met5D | TTTATTCTTCACCTCGTTTTCATTC |
| met10A | GCAGTAGTTCTCGAGACCACC |
| met10D | TTAGTATAATGTGATGGTTAGTTTT |
| met17A | AGGGTTCGAATCCCTTAGC |
| met17D | ACAGCTTTATAGGAGGCGT |

### Supplementary Table 2

| YPGal-Control | | | |
|---|---|---|---|
| Accession | Protein Name | Unused Score | Coverage (%) |
| ACAC_YEAST | Acetyl-CoA carboxylase | 59.56 | 44.42 |
| PYC1_YEAST | Pyruvate carboxylase 1 | 30.46 | 46.10 |
| SYEC_YEAST | Glutamate--tRNA ligase cytoplasmic | 16.73 | 47.18 |
| ARC1_YEAST | tRNA-aminoacylation cofactor ARC1 | 10.27 | 39.10 |
| RS17B_YEAST | 40S ribosomal protein S17-B | 8.00 | 75.00 |
| PYC2_YEAST | Pyruvate carboxylase 2 | 2.00 | 42.03 |
| YJ41B_YEAST | Transposon Ty4-J Gag-Pol polyprotein | 2.00 | 16.75 |
| RS25B_YEAST | 40S ribosomal protein S25-B | 2.00 | 55.56 |

| YPGal- SSH | | | |
|---|---|---|---|
| Accession | Protein Name | Unused Score | Coverage (%) |
| ARC1_YEAST | tRNA-aminoacylation cofactor ARC1 | 18.28 | 37.23 |
| ADH1_YEAST | Alcohol dehydrogenase 1 | 14.00 | 27.01 |
| ALF_YEAST | Fructose-bisphosphate aldolase | 12.01 | 30.36 |
| EF1A_YEAST | Elongation factor 1-alpha | 11.53 | 29.69 |
| ENO2_YEAST | Enolase 2 | 10.63 | 47.60 |
| HSP71_YEAST | Heat shock protein SSA1 | 10.20 | 40.81 |
| PGK_YEAST | Phosphoglycerate kinase | 8.02 | 37.50 |
| KPYK1_YEAST | Pyruvate kinase 1 | 8.01 | 40.00 |
| RL27B_YEAST | 60S ribosomal protein L27-B | 4.54 | 38.24 |
| RS3A1_YEAST | 40S ribosomal protein S1-A | 4.48 | 38.04 |
| ACAC_YEAST | Acetyl-CoA carboxylase | 4.38 | 15.63 |
| RS8B_YEAST | 40S ribosomal protein S8-B | 4.15 | 38.99 |
| TPIS_YEAST | Triosephosphate isomerase | 4.13 | 29.44 |
| RL10_YEAST | 60S ribosomal protein L10 | 4.00 | 53.39 |
| GAL1_YEAST | Galactokinase | 4.00 | 16.67 |
| G3P3_YEAST | Glyceraldehyde-3-phosphate dehydrogenase 3 | 4.00 | 28.92 |
| PDC1_YEAST | Pyruvate decarboxylase isozyme 1 | 4.00 | 11.90 |
| PMG1_YEAST | Phosphoglycerate mutase 1 | 4.00 | 27.13 |
| RL3_YEAST | 60S ribosomal protein L3 | 2.00 | 39.02 |
| VDAC1_YEAST | Mitochondrial outer membrane protein porin 1 | 2.00 | 32.86 |
| RS4B_YEAST | 40S ribosomal protein S4-B | 2.00 | 19.92 |
| RL6A_YEAST | 60S ribosomal protein L6-A | 2.00 | 26.70 |
| HSP74_YEAST | Heat shock protein SSA4 | 1.46 | 33.02 |

### Supplementary Table 2

| YPD-Control | | | |
|---|---|---|---|
| Accession | Protein Name | Unused Score | Coverage (%) |
| ACAC_YEAST | Acetyl-CoA carboxylase | 132.24 | 70.67 |
| PYC1_YEAST | Pyruvate carboxylase 1 | 48.95 | 70.08 |
| SYEC_YEAST | Glutamate--tRNA ligase cytoplasmic | 47.71 | 70.20 |
| ARC1_YEAST | tRNA-aminoacylation cofactor ARC1 | 31.70 | 76.60 |
| DUR1_YEAST | Urea amidolyase | 21.94 | 54.55 |
| RS17B_YEAST | 40S ribosomal protein S17-B | 14.00 | 95.59 |
| COPA_YEAST | Coatomer subunit alpha | 11.51 | 46.63 |
| PYC1_YEAST | Pyruvate carboxylase 1 | 6.35 | 56.53 |
| RL37A_YEAST | 60S ribosomal protein L37-A | 4.18 | 86.36 |
| RL17A_YEAST | 60S ribosomal protein L17-A | 4.07 | 81.52 |
| RL37B_YEAST | 60S ribosomal protein L37-B | 4.01 | 87.50 |
| RS16B_YEAST | 40S ribosomal protein S16-B | 3.21 | 94.41 |
| RL35B_YEAST | 60S ribosomal protein L35-B | 2.34 | 71.67 |
| RL43B_YEAST | 60S ribosomal protein L43-B | 2.13 | 69.57 |
| RS30B_YEAST | 40S ribosomal protein S30-B | 2.01 | 87.30 |
| RS15_YEAST | 40S ribosomal protein S15 | 2.00 | 57.04 |
| RS19B_YEAST | 40S ribosomal protein S19-B | 2.00 | 66.67 |
| IRC5_YEAST | Uncharacterized ATP-dependent helicase IRC5 | 1.96 | 32.94 |
| RGT1_YEAST | Glucose transport transcription regulator RGT1 | 1.42 | 18.38 |
| RL13B_YEAST | 60S ribosomal protein L13-B | 1.33 | 80.40 |
| **YPD-SSH** | | | |
| Accession | Protein Name | Unused Score | Coverage (%) |
| ARC1_YEAST | tRNA-aminoacylation cofactor ARC1 | 26.13 | 67.02 |
| KPYK1_YEAST | Pyruvate kinase 1 | 24.17 | 63.40 |
| EF1A_YEAST | Elongation factor 1-alpha | 24.03 | 55.90 |
| ADH1_YEAST | Alcohol dehydrogenase 1 | 18.61 | 54.60 |
| ACAC_YEAST | Acetyl-CoA carboxylase | 18.51 | 43.48 |
| ALF_YEAST | Fructose-bisphosphate aldolase | 14.03 | 37.88 |
| ENO2_YEAST | Enolase 2 | 12.09 | 55.38 |
| RS11B_YEAST | 40S ribosomal protein S11-B | 11.05 | 83.33 |
| HSP71_YEAST | Heat shock protein SSA1 | 10.59 | 32.87 |
| G3P3_YEAST | Glyceraldehyde-3-phosphate dehydrogenase 3 | 8.06 | 49.10 |
| PGK_YEAST | Phosphoglycerate kinase | 8.00 | 54.09 |
| HSP75_YEAST | Heat shock protein SSB1 | 6.58 | 32.79 |
| RL3_YEAST | 60S ribosomal protein L3 | 6.15 | 60.98 |
| RL27B_YEAST | 60S ribosomal protein L27-B | 6.10 | 76.47 |
| RS8B_YEAST | 40S ribosomal protein S8-B | 6.09 | 53.50 |
| RS16B_YEAST | 40S ribosomal protein S16-B | 6.03 | 70.63 |
| PDC1_YEAST | Pyruvate decarboxylase isozyme 1 | 6.00 | 22.56 |
| PST2_YEAST | Protoplast secreted protein 2 | 6.00 | 36.36 |
| RL20B_YEAST | 60S ribosomal protein L20-B | 5.70 | 56.40 |
| RL6B_YEAST | 60S ribosomal protein L6-B | 4.80 | 38.07 |
| RS3_YEAST | 40S ribosomal protein S3 | 4.70 | 63.33 |
| RS4B_YEAST | 40S ribosomal protein S4- | 4.63 | 60.54 |
| EF2_YEAST | Elongation factor 2 | 4.48 | 31.71 |
| RL10_YEAST | 60S ribosomal protein L10 | 4.11 | 44.80 |
| RS3A2_YEAST | 40S ribosomal protein S1-B | 4.09 | 52.16 |

Supplementary Table 2

| | | | |
|---|---|---|---|
| TKT1_YEAST | Transketolase 1 | 4.03 | 31.32 |
| RL4A_YEAST | 60S ribosomal protein L4-A | 4.00 | 67.68 |
| PMG1_YEAST | Phosphoglycerate mutase 1 | 4.00 | 54.25 |
| RS20_YEAST | 40S ribosomal protein S20 | 4.00 | 71.07 |
| TPIS_YEAST | Triosephosphate isomerase | 4.00 | 25.81 |
| CYPH_YEAST | Peptidyl-prolyl cis-trans isomerase | 4.00 | 25.31 |
| RL7A_YEAST | 60S ribosomal protein L7-A | 3.50 | 72.54 |
| RL13B_YEAST | 60S ribosomal protein L13-B | 2.89 | 70.35 |
| RL15B_YEAST | 60S ribosomal protein L15-B | 2.86 | 76.47 |
| PYR1_YEAST | Protein URA2 | 2.50 | 21.54 |
| RL30_YEAST | 60S ribosomal protein L30 | 2.16 | 50.48 |
| MPG1_YEAST | Mannose-1-phosphate guanyltransferase | 2.11 | 30.75 |
| RS24B_YEAST | 40S ribosomal protein S24-B | 2.05 | 68.15 |
| RS19B_YEAST | 40S ribosomal protein S19-B | 2.01 | 64.44 |
| RL6A_YEAST | 60S ribosomal protein L6-A | 2.00 | 47.16 |
| CG13_YEAST | G1/S-specific cyclin CLN3 | 2.00 | 18.79 |
| YCP4_YEAST | Flavoprotein-like protein YCP4 | 2.00 | 34.01 |
| HAL9_YEAST | Halotolerance protein 9 | 1.34 | 13.20 |

Supplementary Table 3

### Protein activity values

**GAPDH 2 h**

| NaHS mM | Media ± SD |
|---|---|
| 0 | 0.2364 ± 0.01 |
| 0.1 | 0.3324 ± 0.03 |
| 0.25 | 0.3505 ± 0.02 |

**GAPDH 7 h**

| NaHS mM | Media ± SD |
|---|---|
| 0 | 0.2832 ± 0.05 |
| 0.1 | 0.3716 ± 0.02 |
| 0.25 | 0.2837 ± 0.02 |

**PK 2 h**

| NaHS mM | Media ± SD |
|---|---|
| 0 | 0.1216 ± 0.02 |
| 0.1 | 0.2909 ± 0.02 |
| 0.25 | 0.3398 ± 0.09 |

**PK 7 h**

| NaHS mM | Media ± SD |
|---|---|
| 0 | 0.3686 ± 0.02 |
| 0.1 | 0.4577 ± 0.06 |
| 0.25 | 0.3286 ± 0.05 |
